# Cold tolerance in rice plants is partially controlled by root responses

**DOI:** 10.1101/2020.01.25.919464

**Authors:** Angie Geraldine Sierra Rativa, Artur Teixeira de Araújo Junior, Daniele da Silva Friedrich, Rodrigo Gastmann, Thainá Inês Lamb, Igor de Vargas, Alexsander dos Santos Silva, Ândrea Pozzebon-Silva, Janete Mariza Adamski, Janette Palma Fett, Felipe Klein Ricachenevsky, Raul Antonio Sperotto

## Abstract

Rice (*Oryza sativa* L.) ssp. *indica* is the most cultivated species in the South of Brazil. However, these plants face low temperature stress from September to November, which is the period of early sowing, affecting plant development during the initial stages of growth, and reducing rice productivity. This study aimed to characterize the root response to low temperature stress during the early vegetative stage of two rice genotypes contrasting in their cold tolerance (CT, cold-tolerant; and CS, cold-sensitive). Root dry weight and length, as well as number of root hairs, were higher in CT than CS when exposed to cold treatment. Histochemical analyses indicated that roots of CS genotype present higher levels of lipid peroxidation and H_2_O_2_ accumulation, along with lower levels of plasma membrane integrity than CT under low temperature stress. RNAseq analyses revealed that the contrasting genotypes present completely different molecular responses to cold stress. The number of over-represented functional categories was lower in CT than CS under cold condition, suggesting that CS genotype is more impacted by low temperature stress than CT. Several genes might contribute to rice cold tolerance, including the ones related with cell wall remodeling, cytoskeleton and growth, signaling, antioxidant system, lipid metabolism, and stress response. On the other hand, high expression of the genes *SRC2* (defense), *root architecture associated 1* (growth), *ACC oxidase*, *ethylene-responsive transcription factor*, and *cytokinin-O-glucosyltransferase 2* (hormone-related) seems to be related with cold sensibility. Since these two genotypes have a similar genetic background (sister lines), the differentially expressed genes found here can be considered candidate genes for cold tolerance and could be used in future biotechnological approaches aiming to increase rice tolerance to low temperature.

## Introduction

Rice (*Oryza sativa* L.) is one of the most important cereal crops, providing food for more than half of the world’s population (Zhang et al. 2017). Rice production can be seriously affected by environmental stresses, including low temperature, since it is normally grown in tropical and temperate climate zones, and therefore is sensitive to chilling (Mukhopadhyay et al. 2004; Byun et al. 2017). Cold can be harmful from germination to reproductive stages, in most cases resulting in decreased crop yield (Ma et al. 2009; Cruz et al. 2013). Therefore, cold-tolerant plants are able to grow better than cold-sensitive ones under low temperature conditions (Cabello et al. 2014).

Given its tropical origin, cultivated rice germplasm genetic variability is very limited to identify cold stress tolerance genes and alleles (Byun et al. 2017). A few examples of alleles associated with low temperature tolerance are derived from *japonica* subspecies, as *COLD1^jap^* and *bZIP73^jap^* (Ma et al. 2015; Liu et al. 2018). Indeed, transferring the *japonica* alleles to *indica* rice genotypes increased its cold tolerance. Even though *japonica* subspecies is more adapted to temperate climates (Cruz et al. 2013), few *indica* rice cultivars can tolerate stressful conditions and are able to grow under low temperature (Sanghera et al. 2011).

Our group previously identified two *indica* rice genotypes (sister lines with similar genetic background) with contrasting levels of cold tolerance during germination and early vegetative stage (Dametto et al. 2015; Adamski et al. 2016; Sperotto et al. 2018). Parallel physiological and transcriptomic analyses under cold treatment (13°C/ 7 days and 10°C/ 6 hours for germination and vegetative stage, respectively) indicated protective and more active processes in the cold-tolerant genotype when compared to the cold-sensitive one, including cell division and expansion processes, membrane fatty acid unsaturation, antioxidant capacity, cellulose deposition in the cell wall, photosynthetic performance under cold stress/ photosynthetic performance recovery after cold stress, and expression of numerous genes with diverse functions related to chilling stress response, including *Temperature-induced lipocalin-2* and transcription factors *SNAC1* and *OsERD15*.

Optimal root development and distribution play multiple roles in rice growth, including anchorage of the plant and acquisition of water and nutrient elements, which are heterogeneously distributed in soil (Gowda et al. 2011; Lynch 2013). Although the understanding about rice roots has been expanded in the last decades, and a wide natural variation of root system architecture has been reported (Uga et al. 2009), much remains to be investigated regarding root morphology and physiology, particularly in root genetics (Wu and Cheng, 2014), mostly because there is a close relation between above ground traits (including grain yield and quality) and underground roots (Yang, 2011; Ahamed et al. 2012; Neilson et al. 2013). Information on gene networks involved in root formation has been accumulated for *Arabidopsis thaliana*, but our knowledge of these aspects in rice is still limited (Rebouillat et al. 2009; Coudert et al. 2010). As roots can be considered the foundation of rice development, dissecting genetic and molecular mechanisms controlling rice root responses is critical for the development of new rice genotypes that are better adapted to adverse conditions (Rebouillat et al. 2009; Coudert et al. 2010; Wu and Cheng, 2014), including cold stress. Unfortunately, only a few studies of cold tolerance in rice have taken root responses into account (Hashimoto et al. 2009; Lee et al. 2009; Ahamed et al. 2012; Neilson et al. 2013; Xiao et al. 2014; Yang et al. 2015; Zhu et al. 2015).

In our previous works of rice genotypes with contrasting levels of cold-tolerance (cited above), we repeatedly noted that roots of the cold-tolerant and cold-sensitive genotypes seemed different under low temperature stress, being longer and thicker in cold-tolerant plants. For this reason, we decided to better understand the root responses of these contrasting rice genotypes to cold stress, and how that could be related to cold tolerance. In order to identify and characterize novel genes and processes involved with rice root cold tolerance, we performed parallel transcriptomic (10°C for 24 h) and physiological analyses. Our findings could be useful for future biotechnological and breeding efforts aiming to enhance cold tolerance in rice plants.

## Material and methods

### Plant materials and cold treatment

Previously, we identified low temperature tolerant (IRGA 959-1-2-2F-4-1-4-A) and sensitive (IRGA 959-1-2-2F-4-1-4-D-1-CA-1) sister lines from the *indica* subspecies (Dametto et al. 2015; Adamski et al. 2016; Sperotto et al. 2018). Both genotypes were germinated for five days in distilled water and then transferred to holders positioned over plastic pots covered with aluminum foil and containing 300 mL of nutrient solution (as described by Ricachenevsky et al. 2011). The pH of the nutrient solution was adjusted to 5.4. Plants were kept at 28°C ± 1°C under photoperiod of 16 h /8 h light/dark (150 μmol.m^-2^.s^-1^). Solutions were replaced every two days. At the three-leaf stage (approximately 30 days), plants were transferred to a growth chamber and maintained at 10°C (or control condition at 28°C) for up to 10 days (light intensity of 100 μmol.m^-2^.s^-1^).

### Root measurements

Roots (n = 16) from plants submitted to control and cold treatments were measured and oven-dried at 65°C until constant weight, in order to obtain the length (cm) and dry weight (mg), respectively. Root diameter (mm) was measured 1 cm from the apex, using ImageJ software (n = 15). Root hair density was measured in 1 cm sections (always derived from the same location, i.e. 1 cm from the apex) of roots from plants submitted to control and cold treatments (n = 16).

### In situ histochemical localization of H_2_O_2_

*In situ* accumulation of H_2_O_2_ was detected by histochemical staining with diaminobenzidine (DAB), according to Shi et al. (2010), with minor modifications. Roots were excised and immersed in a 1 mg ml^-1^ solution of DAB (pH 3.8) in 10 mM phosphate buffer (pH 7.8) at room temperature. Immersed roots were illuminated for 8 h until brown spots were visible, which were derived from the reaction of DAB with H_2_O_2_. Roots were kept in 70% ethanol for taking pictures with a digital camera coupled to a stereomicroscope.

### Detection of cell death by loss of plasma membrane integrity

To determine changes in viability of cells by cold treatment, excised roots (n = 5 per genotype/ treatment) were immersed for 5 h in a 0.25% (w/v) aqueous solution of Evans Blue (Romero-Puertas et al. 2004). Roots were photo documented with a digital camera coupled to a stereomicroscope.

### Evaluation of lipid peroxidation

Histochemical detection of lipid peroxidation was performed as described by Pompella et al. (1987). In brief, roots (n = 5 per genotype/ treatment) were stained with Schiff’s reagent for 60 min, to detect aldehydes originating from lipid peroxidation. After the reaction with Schiff’s reagent, roots were rinsed with a sulfite solution (0.5% [w/v] K_2_S_2_O_5_ in 0.05 M HCl). The stained roots were kept in the sulfite solution to retain the staining color. The roots stained with Schiff’s reagent were photo documented with a digital camera coupled to a stereomicroscope.

### Detection of cellulose deposition

For analyzing the cellulose deposition in root cell walls of the CT and CS genotypes, 30-day old plants (n = 10 per genotype/ treatment) were submitted to control and cold conditions for 72 h. Root samples were washed and then fixed in a glutaraldehyde 1% + formaldehyde 4% solution in 0.1 M phosphate buffer (pH 7.2), for 24 h, and subsequently went through a dehydration process through an alcoholic series, starting at 10% up to 70% for 20 min, and maintained in 70% ethanol until cutting. Root transversal sections were obtained 1 cm from the apex (3-4 sections per root) using freehand technique. The cuts were hydrated and stained in Congo red 1% for 1 min, which colors cellulose-rich cells (Zheng et al. 2018). After staining, the sections were washed with MilliQ water until dye removal. The colored sections were observed using a Zeis Scope A.1 optical microscope coupled to an Axio CAM 506 color digital camera.

### RNA extraction and comparative transcriptomic profiling by RNAseq

Total RNA was extracted from rice roots using Concert Plant RNA Reagent (Invitrogen^®^) and treated with DNase I (Invitrogen^®^). Approximately 20 μg of total RNA were used in high-throughput cDNA sequencing by Illumina HiSeq 2000 technology (Fasteris SA, Plan-les-Ouates, Switzerland - http://www.fasteris.com/). We constructed two individual single-end cDNA library for each rice genotype (cold tolerant and sensitive), after 24 h of cold or control treatment, totalizing eight libraries (four plants per library). The cDNA libraries were prepared according to Illumina’s protocols. Briefly, RNAseq was performed using the following successive steps: poly-A purification; cDNA synthesis using a poly-T primer, shotgun method to generate inserts of approximately 500 bp; 3p and 5p adapter ligations; pre-amplification; colony generation; and Illumina single-end 50 bp sequencing.

Low quality reads (FASTq value <13) were removed, and 3p and 5p adapter sequences were trimmed using Genome Analyzer Pipeline (Fasteris). The remaining low-quality reads with ‘n’ were removed using Python script. After cleaning the data (low quality reads, adapter sequences), the mRNAseq data from the eight libraries was aligned to the rice genome using the software Spliced Transcripts Alignment to a Reference (STAR) (Dobin et al. 2013). Only sequences with up to two mismatches to the rice reference genome (http://rice.plantbiology.msu.edu/pub/data/Eukaryotic_Projects/o_sativa/annotation_dbs /pseudomolecules/version_7.0/all.dir/all.seq) were used. The SAM files from STAR were then processed using Python scripts to assign the frequencies of each read and map them onto references. For data normalization, we used the scaling normalization method (Robinson et al. 2010). To assess whether the sequences were differentially expressed, we used the R package EdgeR (Robinson et al. 2010). We considered that the sequences were differentially expressed if they had an adjusted p-value < 0.05 and at least 2-fold change. The RNA-seq data obtained in this work was deposited on the Gene Expression Omnibus database (GEO - https://www.ncbi.nlm.nih.gov/geo/ - Barrett et al. 2013) and can be accessed using the code

### Gene Ontology (GO) terms enrichment analysis

The *loci* from differentially expressed genes with increased expression in plants from the cold-tolerant or cold-sensitive genotypes were used to search for enriched Gene Ontology (GO) terms comparing both datasets. Enrichment analysis was performed using Plant GeneSet Enrichment Analysis Toolkit (PlantGSEA - http://structuralbiology.cau.edu.cn/PlantGSEA/ - Yi et al. 2013) built-in Fisher’s Exact Test. The significance threshold of gene set enrichments was false discovery rate (FDR) < 0.05. From a total of 202 (control condition) and 582 (cold condition) differentially expressed genes, 200 and 569 could be assigned a GO term, respectively, and were thus considered in the GO enrichment analysis.

### Gene expression analysis by quantitative RT-PCR

To confirm the high-quality of deep sequencing results, RT-qPCR was used to check the gene expression of ten putative cold tolerance-related genes. Total RNA was extracted from rice roots of plants exposed to 10°C for 24 h using Concert Plant RNA Reagent (Invitrogen) and treated with DNase I (Invitrogen). First-strand cDNA synthesis was performed with reverse transcriptase (M-MLV, Invitrogen) using 1 μg of RNA. RT-qPCRs were carried out in a StepOne Real-Time Cycler (Applied Biosystems). All primers (listed in Supplementary Table 1) were designed to amplify 100-150 bp of the 3’-UTR of the genes and to have similar Tm values (60 ± 2°C). Reaction settings were composed of an initial denaturation step of 5 min at 94°C, followed by 40 cycles of 10 s at 94°C, 15 s at 60°C (fluorescence data collection) and 15 s at 72°C; samples were held for 2 min at 40°C for annealing of the amplified products and then heated from 55 to 99°C with a ramp of 0.1°C /s to produce the denaturing curve of the amplified products. RT-qPCRs were carried out in 20 μl final volume composed of 10 μl of each reverse transcription sample diluted 100 times, 2 μl of 10X PCR buffer, 1.2 μl of 50 mM MgCl2, 0.1 μl of 5 mM dNTPs, 0.4 μl of 10 μM primer pairs, 4.25 μl of water, 2.0 μl of SYBR green (1:10,000, Molecular Probe), and 0.05 μl of Platinum Taq DNA Polymerase (5 U/μl, Invitrogen, Carlsbad, CA, USA). Gene expression was evaluated using a modified 2^-ΔCT^ method (Schmittgen and Livak 2008), which takes into account the PCR efficiencies of each primer pair (Relative expression TESTED GENE / CONTROL GENE = (PCReff CG)^Ct CG^ / (PCReff TG)^Ct TG^). Rice Ubiquitin (*OsUBQ5*) was used as the control gene. Experiments were performed with three biological and four technical replicates, collected in a second experiment (i.e. distinct from the one for RNAseq).

### Statistical analysis

When appropriate, means were statistically compared by the Student’s *t*-test (*p*-value ≤ 0.05, 0.01 and 0.001) using SPSS Base 21.0 for Windows (SPSS Inc., USA).

## Results

### Physiological characterization of cold-tolerant and cold-sensitive roots

Thirty-day old plants were submitted to control (28°C) or cold (10°C) condition for up to 10 days. As seen in Figure 1, root dry weight of both cold-tolerant (CT) and cold-sensitive (CS) plants decreased in cold conditions. Also, root dry weight of CT plants under cold condition was higher than CS ones during the entire evaluated period. Under control condition we detected higher variation, even so the root dry weight was higher in CT than CS plants in most of the evaluated periods. Root length and root hair density presented higher values in CT plants than in CS ones submitted to cold stress, while no difference was detected under control condition (Figure 1). Histochemical analyses showed that CS roots present high levels of lipid peroxidation (Figure 2a), loss of plasma membrane integrity (indicative of cell death - Figure 2b), and H_2_O_2_ accumulation (Figure 2c), suggesting that roots from CT plants are more efficiently protected of and less damaged by the oxidative stress caused by low temperature than roots from CS plants. Such differences in physiological responses, together with the fact that most cold-tolerance studies are performed only with rice leaves, encouraged us to better understand the molecular responses of these roots under low temperature stress using RNAseq.

**Figure 1.**
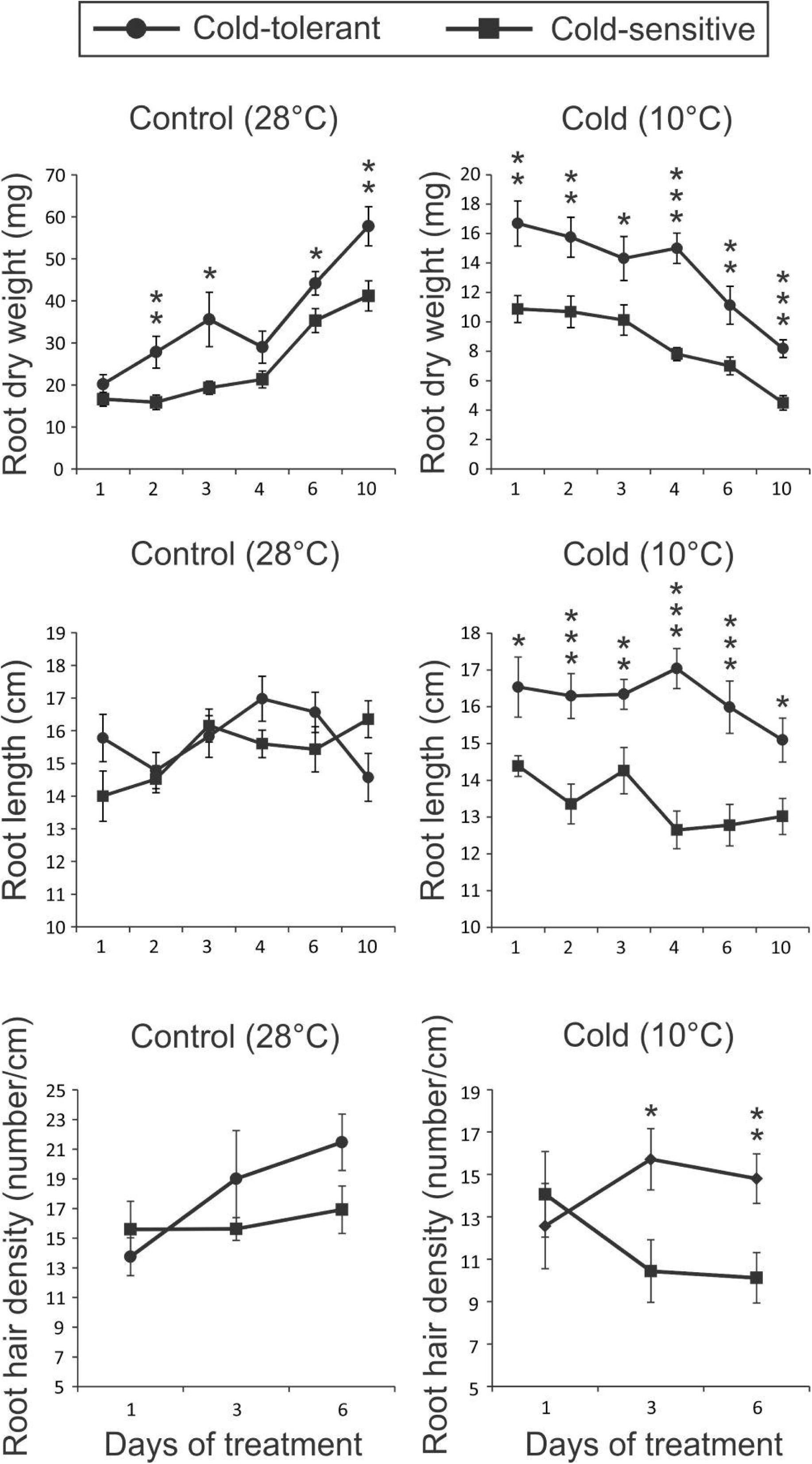
Root dry weight, length, and hair density of thirty-day old cold-tolerant and cold-sensitive plants under control (28°C) or cold (10°C) condition. Represented values are the averages of sixteen samples ± SE. Mean values with one, two or three asterisks are different by Student’s *t* test (*p*-value ≤ 0.05, 0.01 or 0.001, respectively).

**Figure 2.**
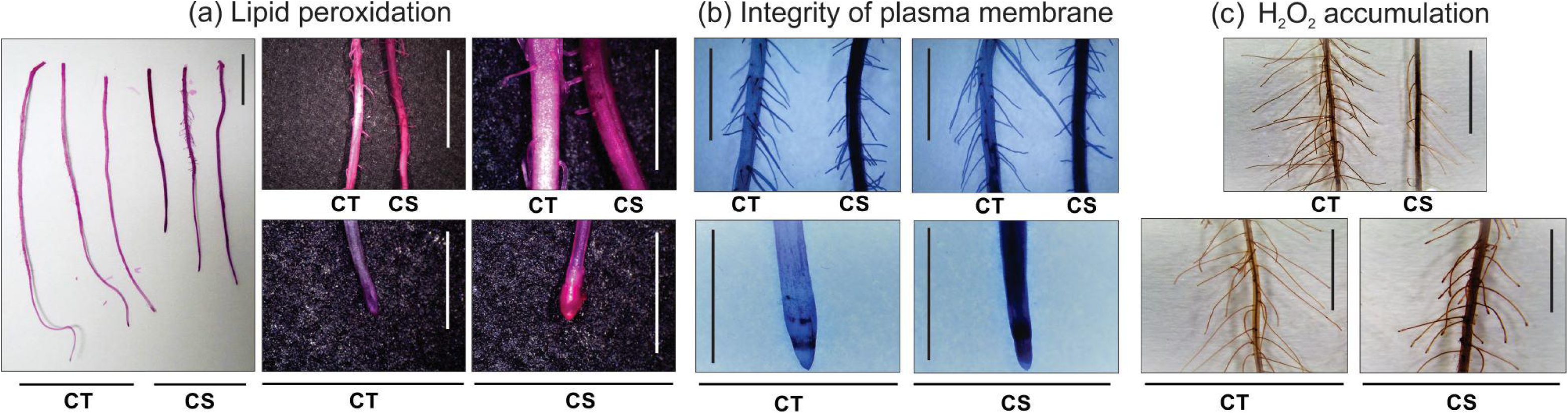
Root histochemical analysis. Lipid peroxidation (a), loss of plasma membrane integrity (b), and H_2_O_2_ accumulation (c) by Schiff, Evans Blue, and DAB reagents, respectively, in thirty-day old roots of cold-tolerant (CT) and cold-sensitive (CS) plants after ten (a) or three days (b and c) under cold (10°C) condition. The positive staining (detected in higher levels in cold-sensitive roots) in the photomicrographs shows as bright images (pink-color for Schiff, blue-color for Evans Blue, and brown-color for DAB). Bars in figures indicate 1.0 cm.

### Overview of cold-tolerant and cold-sensitive root cDNA libraries sequencing

Roots from plants of both genotypes (CT and CS) under cold (10°C) and control (28°C) conditions for 24h were used to identify differentially expressed mRNAs using the Illumina Platform. Since we were interested in understanding the molecular basis of tolerance, we focused on the differences between genotypes in the same experimental condition. Considering the two comparisons performed (Control: CT x CS; and Cold: CT x CS), we found 784 differentially expressed sequences, which are all listed in Supplementary Tables 2 and 3. Under control condition, 104 sequences were more expressed in the CT genotype, and 98 in the CS genotype. Under cold condition, 235 sequences were more expressed in the CT genotype, and 347 in the CS one (Figure 3a). As these genotypes are sister lines (Sperotto et al. 2018), and therefore share a similar genetic background, the low number of differentially expressed sequences between CT and CS was expected. Interestingly, 71 and 69 sequences were more expressed in the CT and CS genotypes, respectively, regardless the tested condition (Figure 3b). These sequences with common expression patterns on the CT and CS genotypes under both treatments are listed in Supplementary Table 4. Also, three sequences presented opposite expression pattern in the CT and CS genotypes: LOC_Os09g29490 (*peroxidase precursor*) and LOC_Os07g14740 (*harpin-induced protein 1 domain containing protein*) were more expressed in the CT genotype under control condition, and more expressed in the CS genotype under cold condition. On the other hand, LOC_Os02g17780 (*ent-kaurene synthase*) was more expressed in the CS genotype under control condition, and more expressed in the CT genotype under cold condition (Supplementary Table 4, marked in gray color).

**Figure 3.**
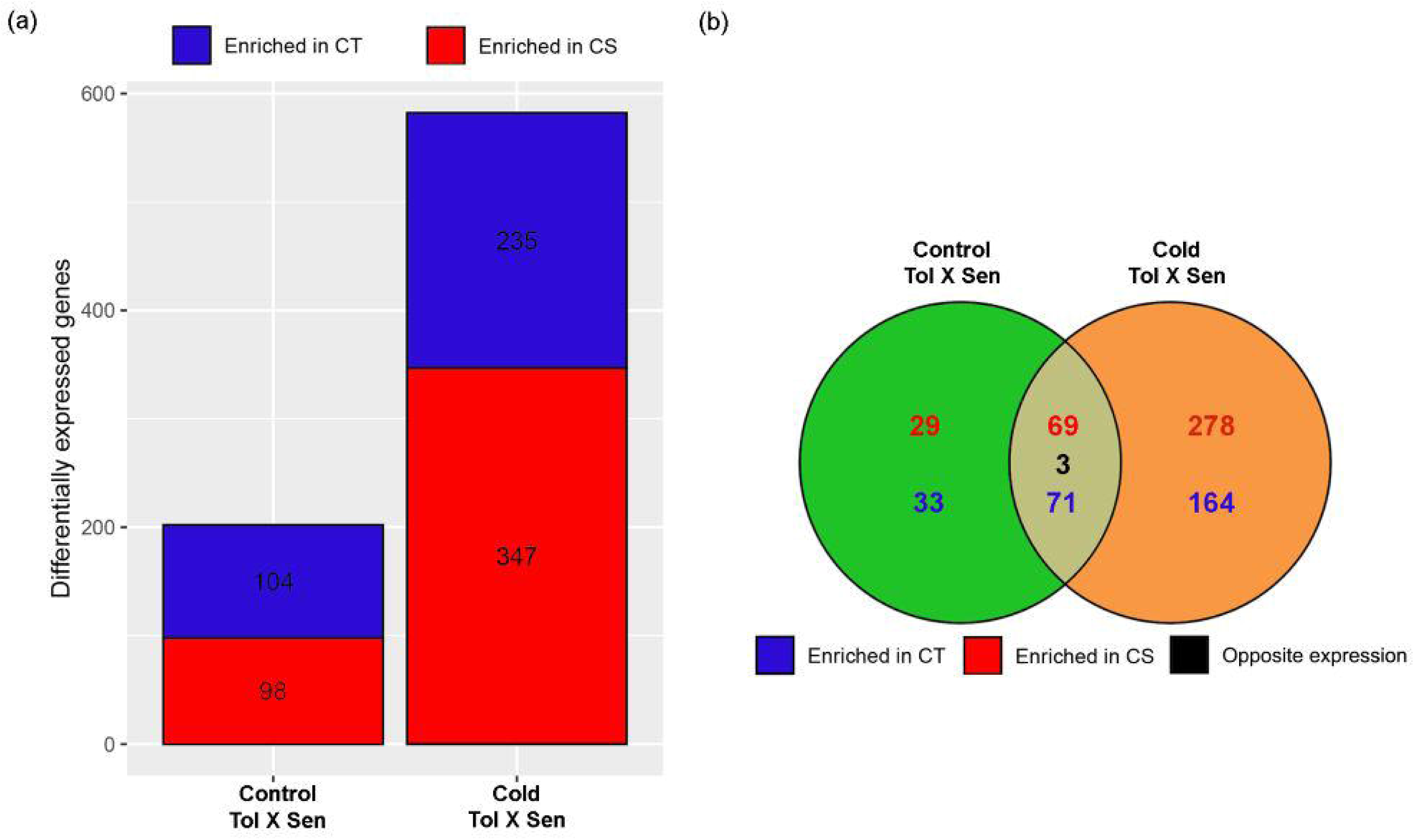
Differentially expressed sequences and similarity between libraries. Number of differentially expressed sequences in different tested conditions (a). Blue bars: enriched in cold-tolerant (CT) plants; Red bars: enriched in cold-sensitive (CS) plants. Venn diagram showing the number of different expressed sequences (specific to each group or common to both analyzed groups) in different conditions (b). Numbers in blue color: enriched in CT plants; numbers in red color: enriched in CS plants; number in black color: number of genes with opposite expression pattern on the two genotypes. Tol = tolerant; Sen = sensitive.

### GO terms and metabolism categories

The GO categories with the highest percentage of annotated genes within each main GO division (biological process, cellular component and molecular function) are shown for both genotypes under control and cold conditions (Supplementary Figure 1). Comparing both genotypes under control condition, GO analysis indicates that the CT genotype presents more active protein metabolic processes (phosphorylation and modification), phosphorus metabolic processes, macromolecule modification, and transferase/kinase activity. On the other hand, other biological processes such as response to stress, defense response, cell death, and apoptotic process, are more active in the CS genotype, along with the molecular function of general binding (Supplementary Figure 1). Under cold treatment, more GO categories were found enriched in both genotypes. Several GO categories were more active in the CT genotype, including general metabolic/biosynthetic processes, cellular component (mostly related to cytoskeleton), protein binding, and synthesis of cellulose. On the other hand, the CS genotype presented more active electron transport and response to stimulus/stress, along with ion binding, kinase activity, and membrane-related processes (Supplementary Figure 1).

It was possible to assign metabolism categories to 55 and 59 genes more expressed in the CT and CS genotype, respectively, after control treatment, and 134 and 223 genes more expressed in the CT and CS genotype, respectively, after cold exposure. It is interesting to note that cold treatment activates several metabolic pathways in both genotypes (Figure 4). Under control condition, protein metabolism- and signaling-related genes are more expressed on the CT genotype, while stress- and secondary metabolism-related genes are more expressed on the CS one (Figure 4a). Under cold condition, most of the metabolic pathways seem more active in the CS genotype (except cell wall- and carbohydrate metabolism-related - Figure 4b), suggesting that the CS genotype is deeply affected by low temperature stress, while the CT genotype presents active cell wall remodeling, probably to reinforce and thicken it. This hypothesis was strengthened by the higher deposition of cellulose in root cells of CT genotype under cold condition when compared with CS (Figure 5a), which probably contributes to the higher root diameter (Figure 5b).

**Figure 4.**
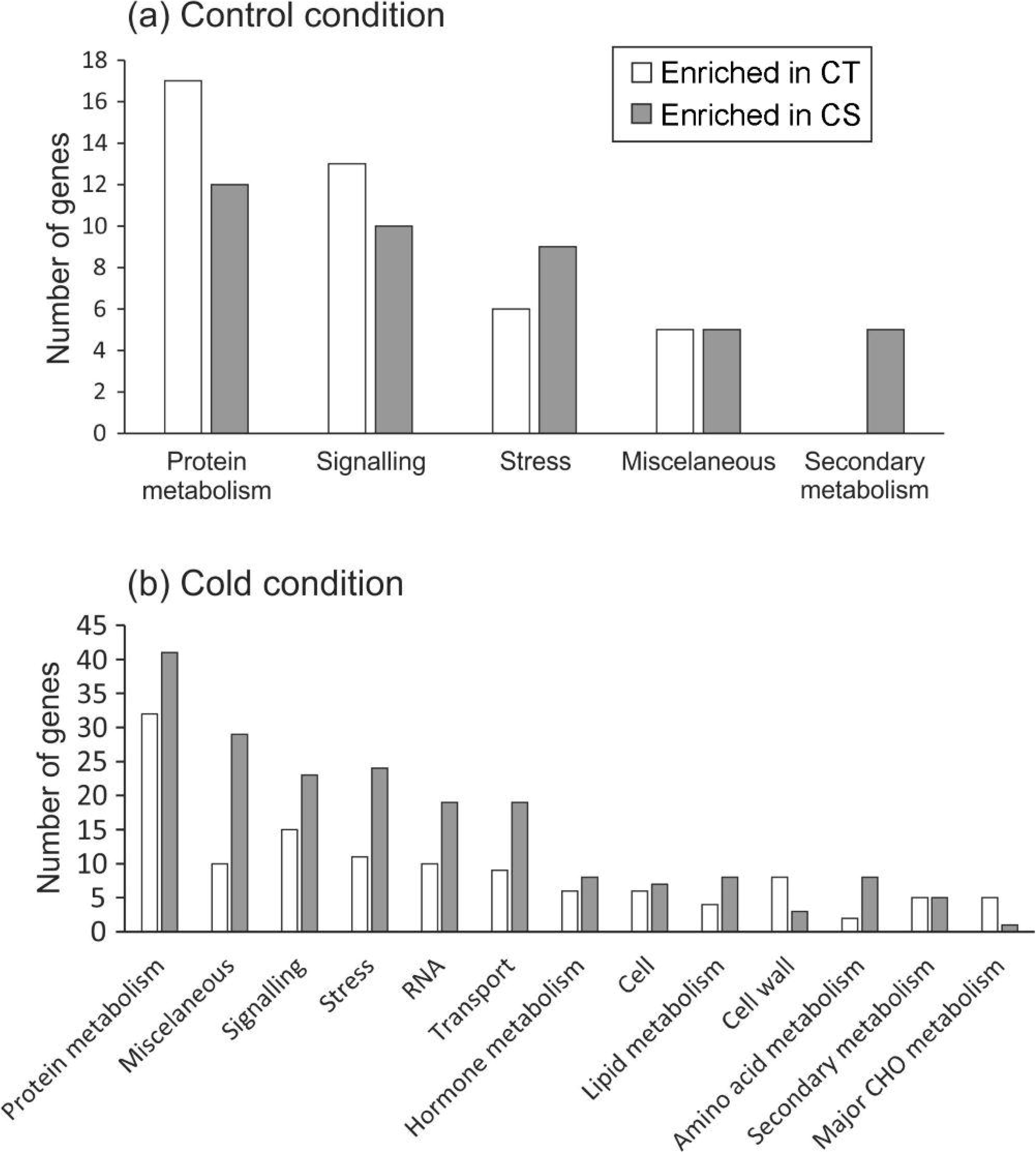
Overview of the MapMan visualization of differences in transcript levels between root transcriptomes from cold-tolerant and cold-sensitive rice genotypes under control (a) and cold (b) conditions. Genes associated with metabolic pathways were analyzed by the MapMan software (http://mapman.gabipd.org/web/guest/mapman).

**Figure 5.**
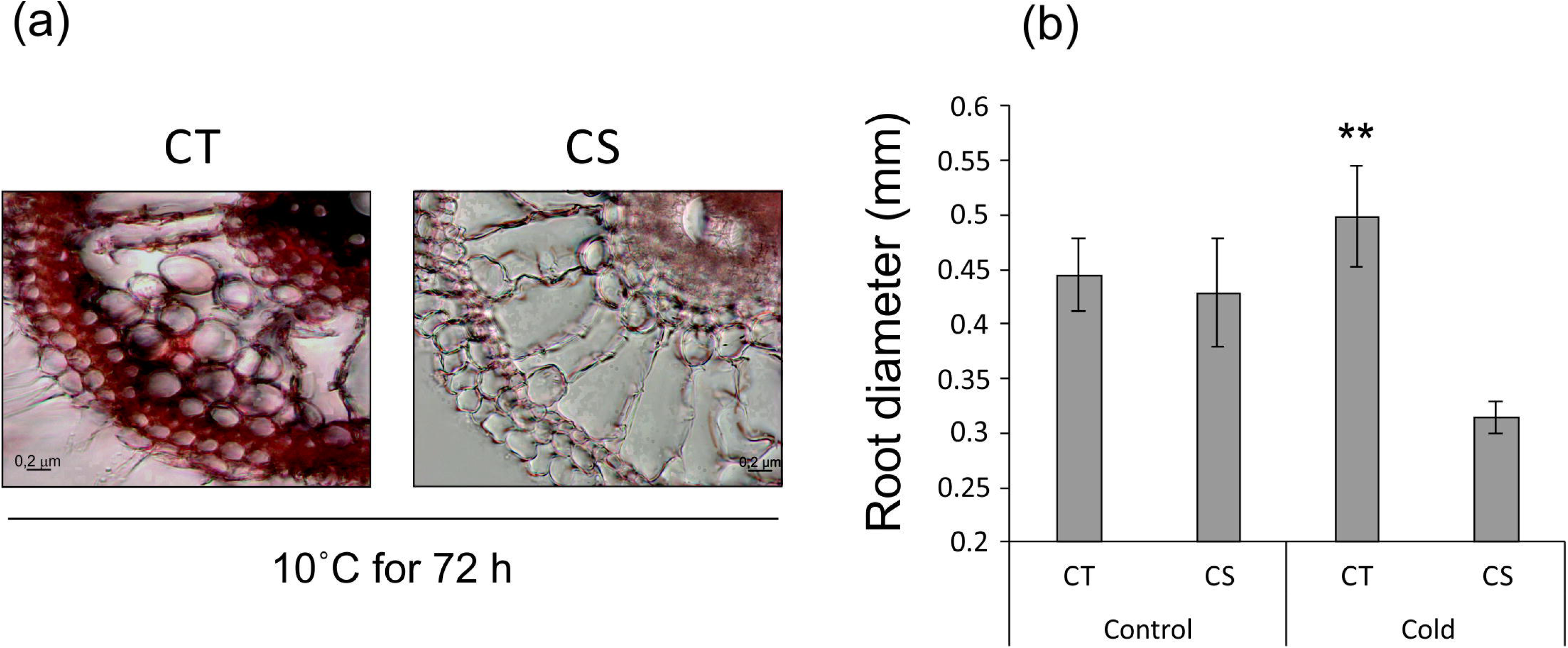
Cellulose staining (a) and root diameter (b) of thirty-day old cold-tolerant (CT) and cold-sensitive (CS) plants under control (28°C) and/or cold (10°C) condition for 72 h. Represented values are the averages of sixteen samples ± SE. Mean values with two asterisks are different by Student’s *t* test (*p*-value ≤ 0.01).

All the 784 differentially expressed genes (DEGs) identified under control or cold conditions were carefully analyzed and, based on the predicted molecular function and available literature, were classified in functional categories (Supplementary Tables 2 and 3). In order to facilitate the overall understanding of the differential gene expression in each functional category presented in Supplementary Tables 2 and 3, we summarize these data in Table 1. Seven functional categories (antioxidant system, carbohydrate metabolism/energy production, cell wall, lipid metabolism, others, signaling, and stress response) were more represented in CT than in CS roots under control conditions, while only five (defense, hormone-related, secondary metabolism, transcription factor, and transport) were more represented in CS than in CT roots (Table 1). Under cold treatment, we detected four different patterns, based on the number of differentially expressed sequences and expression level: 1) functional categories with similar representation in both CT and CS genotypes (carbohydrate metabolism and energy production, lipid metabolism, nucleotide metabolism, secondary metabolism, stress response, and others); 2) functional categories more represented in CT than CS genotype (cell wall, growth, and signaling); 3) functional categories uniquely represented in only one of the genotypes (cytoskeleton in CT, and DNA structure maintenance + photosynthesis in CS); and 4) the most common pattern, containing functional categories more represented in CS than CT genotype (amino acid metabolism, antioxidant system, calcium-related, defense, hormone-related, protein degradation/modification, transcription factor, translation-related, transport, and unknown). It is interesting to highlight the higher expression of cell wall-related genes under control condition and the higher expression of cell wall/growth-related genes under cold condition on the CT genotype (Table 1), which agree with the higher root biomass, length and hair density of this genotype when compared with CS, especially under cold condition (Figure 1). The high number of functional categories more represented in CS than CT genotype (Table 1) agrees with the metabolic pathways data presented in Figure 4b. Several genes belonging to these functional categories are discussed later.

**Table 1.** Schematic representation of the functional categories differently represented in roots of cold-tolerant (CT) or cold-sensitive (CS) rice plants under control and cold conditions. Different shades of gray color represent the four different patterns of response. Number of arrows (one or two) relates to the representation level, which is based on the number of differentially expressed sequences and gene expression level.

| Functional categories | Control condition |  | Cold condition |  |
| --- | --- | --- | --- | --- |
|  | CT | CS | CT | CS |
| Amino acid metabolism | - | - | ↑ | ↑↑ |
| Antioxidant system | ↑↑ | ↑ | ↑ | ↑↑ |
| Calcium-related | - | - | ↑ | ↑↑ |
| Carbohydrate metabolism/energy production | ↑ | - | ↑ | ↑ |
| Cell wall | ↑↑* | ↑ | ↑↑ | ↑ |
| Cytoskeleton | - | - | ↑ | - |
| Defense | ↑ | ↑↑ | ↑ | ↑↑ |
| DNA structure maintenance | - | - | - | ↑ |
| Growth | - | - | ↑↑ | ↑ |
| Hormone-related | ↑ | ↑↑ | ↑ | ↑↑ |
| Lipid metabolism | ↑ | - | ↑ | ↑ |
| Nucleotide metabolism | - | - | ↑ | ↑ |
| Others | ↑↑ | ↑ | ↑ | ↑ |
| Photosynthesis | - | - | - | ↑ |
| Protein degradation/modification | ↑ | ↑ | ↑ | ↑↑ |
| Secondary metabolism | ↑ | ↑↑ | ↑ | ↑ |
| Signaling | ↑↑ | ↑ | ↑↑ | ↑ |
| Stress response | ↑ | - | ↑ | ↑ |
| Transcription factor | ↑ | ↑↑ | ↑ | ↑↑ |
| Translation-related | - | - | ↑ | ↑↑ |
| Transport | - | ↑ | ↑ | ↑↑ |
| Unknown | ↑ | ↑ | ↑ | ↑↑ |

(-): functional category not detected as differentially represented.
(\*): we considered more represented in CT than CS due to the high expression level.

### Confirmation of RNAseq expression patterns using RT-qPCR

To further confirm the quality of our libraries and to evaluate possible induction or repression of expression by the cold treatment, 10 genes were randomly selected for RT-qPCR analyses: six genes with higher expression in the CT plants under both tested conditions, three genes with higher expression on the CS under cold, and one gene with higher expression on the CS under both tested conditions (Supplementary Tables 1, 2 and 3). The differences in expression between the two genotypes after cold exposure, found by the RNAseq method, were confirmed for all tested genes (Figure 6), endorsing the good quality of the datasets.

**Figure 6.**
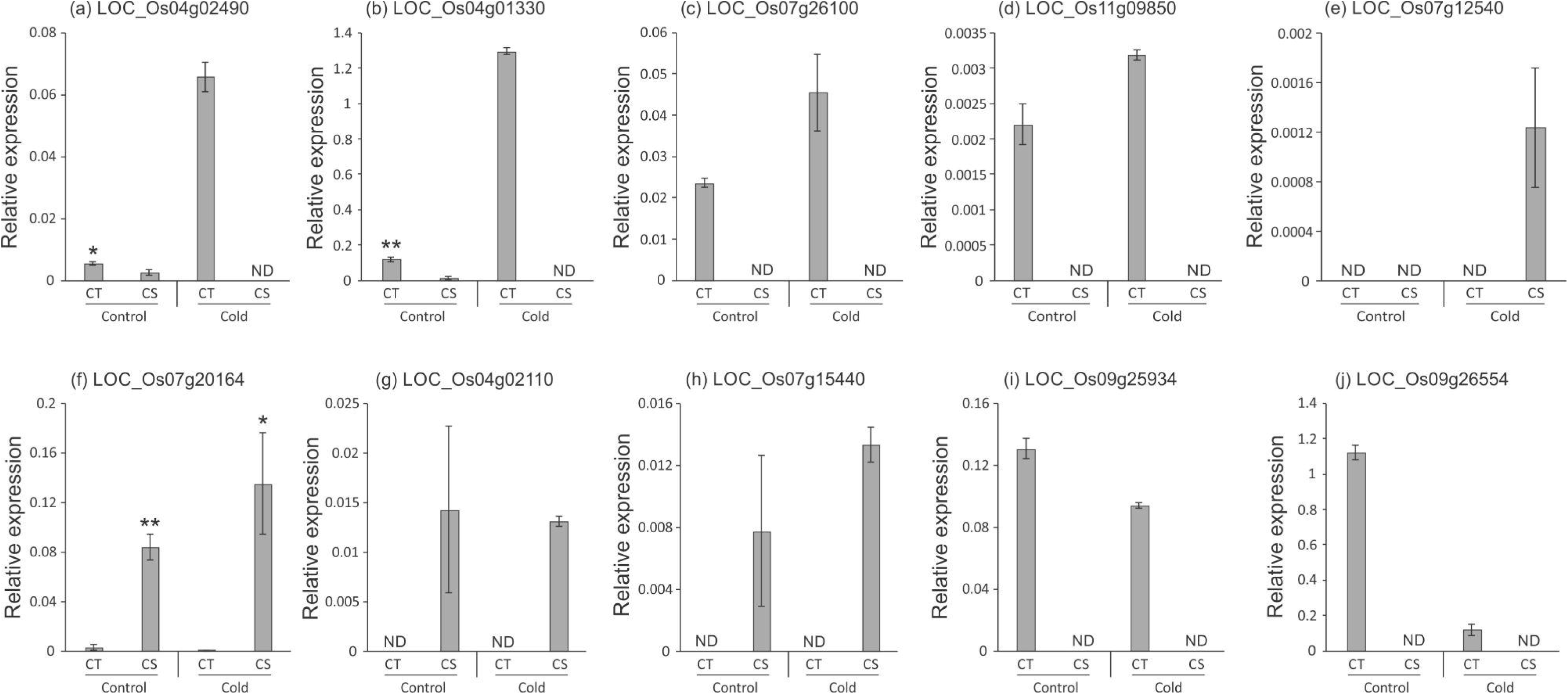
Relative expression levels (RT-qPCR, relative to *OsUBQ5* expression) of selected genes identified by RNAseq in roots of rice plants (CT: cold-tolerant; CS: cold-sensitive) submitted to control or cold conditions: (a) LOC_Os04g02490 (*Expressed protein*), (b) LOC_Os04g01330 (*Expressed protein*), (c) LOC_Os07g26100 (*Expressed protein*), (d) LOC_Os11g09850 (*Expressed protein*), (e) LOC_Os07g12540 (*Expressed protein*), (f) LOC_Os07g20164 (*Expressed protein*), (g) LOC_Os04g02110 (*Disease resistance protein RGA3*), (h) LOC_Os07g15440 (*Alanyl-tRNA synthetase family protein*), (i) LOC_Os09g25934 (*Expressed protein*), and (j) LOC_Os09g26554 (*Expressed protein*). Values are the averages of three samples ± SE. Mean values with one or two asterisks are different by Student’s t test (p ≤ 0.05 and 0,01, respectively). ND = not detected.

## Discussion

### Differential expression under control condition

Root system architecture is a complex trait controlled by several genes (Wachsman et al. 2015). Courtois et al. (2009) summarized 675 rice QTLs for 29 root parameters detected in 12 mapping populations. Even though in our analyses the absolute number of differentially expressed genes was similar under control condition (104 more expressed in the CT genotype, and 98 more expressed in the CS - Figure 3 and Supplementary Table 2), the number of over-represented functional categories was higher in CT than CS under control condition (Table 1). According to Janiak et al. (2018), drought tolerance in barley may be attributed to stressed-like expression patterns present before the occurrence of stress. Therefore, the over-representation of functional categories in roots of CT than CS plants under control condition could be representative of a constitutive tolerance mechanism, as previously suggested for cotton (Zheng et al. 2012).

Under control condition, the CT genotype stood out with higher expression of genes related to antioxidant system, cell wall, lipid metabolism, signaling, stress response, and others (Table 1 and Supplementary Table 2). However, for most of these genes, we cannot establish a direct link with the higher root dry mass of CT than CS plants seen under control condition (Figure 1). Root growth is caused by cell division and elongation. Mutant analyses revealed that genes related to cell wall growth, cell expansion, and auxin signaling are involved in the division and elongation of root cells (Kitomi et al. 2018). Even though the CS genotype presented higher expression of hormone-related genes, the CT genotype presented higher expression of *OsIAA2*, an auxin-responsive gene, along with a cell wall-related gene, *glycosyl hydrolase* (Supplementary Table 2). *OsCel9A*, a rice *glycosyl hydrolase* family gene, plays an essential role in regulating auxin-induced lateral root primordia formation (Yoshida et al. 2006). Auxin signaling is important in overall root morphogenesis/elongation (Kitomi et al. 2018). A mutation in a member of the Auxin (Aux)/Indole-3-acetic acid (IAA) gene family, *OsIAA23*, causes defects in postembryonic quiescent center (QC) maintenance due to the disintegration of the root cap and termination of root growth, suggesting the importance of auxin in rice QC maintenance (Ni et al. 2011). Also, plants overexpressing the auxin biosynthesis gene *OsYUCCA*1 have higher auxin content, which leads to more crown roots (Yamamoto et al. 2007). Susuki et al. (2003) reported the first root-hairless mutant in rice, *rh2*, whose absence of root hairs is likely caused by a shortage of endogenous auxin. Also, WUSCHEL-related homeobox 3A (OsWOX3A) was reported to control root hair formation through the regulation of auxin transport genes (Yoo et al. 2013), further suggesting that auxin is required for root hair development. We also detected higher expression of *phospholipase D*, *phytosulfokine receptor*, and *wali7* in CT than CS plants. The phospholipase D family plays an important role in the regulation of cellular processes in plants, including root growth (Liu et al. 2010). Phytosulfokine-α (PSK-α) is a disulfate pentapeptide described as a growth factor. In Arabidopsis, PSK-α induced root growth (mainly by an increase in cell size) in a dose-dependent manner without affecting lateral root density (Kutschmar et al. 2009). Also, *pskr1-3 pskr2-1* mutant seedlings had shorter roots and hypocotyls than the wild type, whereas *35S:PSKR1* or *35S:PSKR2* seedlings were larger (Hartmann et al. 2013). Janiak et al. (2012) reported that a wali7 domain-containing protein accumulates during the initial stage of root hair development in barley. It is important to highlight that this same gene (*wali7*) was also detected more expressed in the CT than CS under cold condition (Supplementary Table 3).

On the other hand, under control condition the CS genotype showed higher expression of genes related to defense, hormone-related, secondary metabolism, transcription factor, and transport (Table 1 and Supplementary Table 2). As previously discussed, there is no direct relation between the expression of most of these genes with lower root growth of CS than CT plants seen under control condition (Figure 1). One of the hormone-related genes detected with higher expression in the CS than in the CT genotype, *2OG-Fe oxygenase*, participates in ethylene biosynthesis (Dalal et al. 2018). Also, the CS genotype presented higher expression level of the genes *brassinosteroid LRR receptor kinase* and *ent-kaurene synthase* than CT. The first one is known to be involved in normal brassinosteroid perception and plant development, including root growth via hormonal crosstalk with ethylene (Clouse, 2011; Lv et al. 2018). The second one encodes the enzyme catalyzing the second step of the gibberellin (GA) biosynthesis pathway (Margis-Pinheiro et al. 2005), also interacting with ethylene to regulate root growth and defense (Saithong et al. 2015; Tezuka et al. 2015). Under control condition, we detected higher expression of genes related to terpene biosynthesis (*cycloartenol synthase*, *phytoene synthase* and *terpene synthase*) in roots of CS than CT (Supplementary Table 2). It was shown that ethylene treatment can up-regulate *cycloartenol synthase* in the perennial medicinal herb *Dioscorea zingiberensis* (Diarra et al. 2013). Also, ethylene regulates carotenoid biosynthesis (which includes phytoene synthase protein) in rice (Yin et al. 2015), and sesquiterpene biosynthesis in citrus, through the activation of a *terpene synthase* gene (*CsTPS1* - Shen et al. 2016).

Therefore, several ethylene-related genes were found more expressed on the roots of CS than CT plants. It is important to highlight that high levels of ethylene can inhibit root growth (Yin et al. 2015; Qin and Huang, 2018). In Arabidopsis, exogenous application of ethylene inhibits the primary root elongation (Ruzicka et al. 2007). Thus, we hypothesize that roots of CS plants present lower growth under control condition (Figure 1) due to higher ethylene levels than CT.

### Differentially expressed genes under cold condition

Under low temperature, the absolute number of differentially expressed genes was lower in CT than CS (235 more expressed on the CT genotype, and 347 more expressed on the CS - Figure 3 and Supplementary Table 3). Also, the number of over-represented functional categories was lower in CT than CS under cold condition (Table 1), suggesting that the CS genotype is more prone to sense and more impacted by low temperature stress than the CT. Under cold treatment, the CT genotype showed higher expression of genes related to only four categories: cell wall, cytoskeleton, growth, and signaling (Table 1 and Supplementary Table 3).

The cell wall senses and broadcasts stress signals to the interior of the cell, by triggering a cascade of reactions leading to plant tolerance/resistance. Therefore, the study of wall-related genes is particularly relevant to understand the metabolic remodeling triggered by plants in response to exogenous stresses (Guerriero et al. 2014). Several genes involved in the formation and remodeling of plant cell wall, including *glycosyl hydrolase*, *cellulose synthase*, *glycosyl transferase*, *wall-associated kinase (WAK)*, and *glycine-rich cell wall structural protein*, were detected with higher expression in the CT than CS under cold condition (Supplementary Table 3). Glycosyl hydrolases have been implicated in physiologically important processes in plants, such as response to stresses, activation of phytohormones, lignification, and cell wall remodeling (Opassiri et al. 2006). Cellulose, hemicelluloses, and pectins are the main structural polysaccharides of the plant cell wall. Enzymes as glycosyl transferases and cellulose synthases are involved in the synthesis of these polysaccharides and other glycans (De Caroli et al. 2014) and have been previously related with cold tolerance in young rice plants (Dametto et al. 2015; Sperotto et al. 2018; Kong et al. 2019). To our knowledge, this is the first time that these cell wall-related genes are detected with higher expression in rice roots of CT plants when compared to CS ones. Biochemical studies demonstrated that WAK proteins (which play important roles in cell expansion) are covalently bound to pectin in the cell wall and serve as physical links between the extracellular matrix and the cytoplasm and as a signaling component between the cell wall and the cytoplasm (Zhang et al. 2005; de Oliveira et al. 2014), interacting with glycine-rich proteins (Park et al. 2001; Giarola et al. 2016).

The maintenance of cellulose synthesis under stress is necessary for proper stress response. Interestingly, a new protein family named Cellulose Synthase Companion (CC) proteins, was shown to be crucial to the stability of the Cellulose Synthase Complex during salt stress in Arabidopsis. CC proteins promotes the reassembly of microtubules, which are important to drive cellulose deposition at the cell wall (Kesten et al. 2019). Interestingly, some genes related with the functional categories cytoskeleton remodeling and growth were detected with higher expression in CT than CS under cold condition, including *microtubule-associated protein 70 (MAP70)*, *kinesin motor domain containing protein*, *growth regulating factor protein*, *auxin-independent growth promoter protein*, and *RopGEF7* (Supplementary Table 3). Microtubules are a component of the plant cytoskeleton composed of heterodimers of αβ–tubulin. They perform several cellular roles, perhaps most notably coordinating the deposition of cellulose microfibrils in the cell wall (Gardiner, 2013). Kinesins, the ATP-driven microtubule (MT)-based motor proteins, have been reported to be involved in many basic processes, including cell division and organ development (Fang et al. 2018). Recently, Xu et al. (2018) showed that a rice class-XIV kinesin (coded by one of the kinesin genes detected in our study - Os04g53760) actively transports microtubules along each other in a unidirectional manner and enters the nucleus in response to cold, stimulating cell elongation. Growth regulating factors are plant-specific transcription factors important for developmental processes, including root development and coordination of growth processes under adverse environmental conditions (Omidbakhshfard et al. 2015). In Arabidopsis, RopGEF7 (a GTPase activator) is required for root meristem maintenance, which is essential for the maintenance of root growth (Chen et al. 2011). AtRopGEF7 is induced transcriptionally by auxin, while its function is required for the maintenance of normal auxin levels in seedling roots, suggesting that RopGEF7 may integrate auxin-derived positional information (Chen et al. 2011). Even though, we were not able to detect differential expression of auxin-related genes in the CT genotype, what could explain the higher expression of *auxin-independent growth promoter protein* in CT than CS (Supplementary Table 3).

Signaling-related genes were found more expressed in the CT than CS under cold condition, including *receptor-like protein kinase* and *Rapid Alkalinization Factor (RALFL21)* (Supplementary Table 3). Receptor-like kinases are a prominent class of surface receptors that regulate many aspects of the plant life cycle, including cell expansion (Yang et al. 2015). FERONIA (FER), a receptor-like kinase, is involved in brassinosteroids and auxin signaling pathways, which have positive roles in cell growth and development. Such activation occurs through the interaction with a Rapid Alkalinization Factor (RALF1) peptide, triggering a series of downstream events related with plant growth and stress response (Li et al. 2018).

Other genes with higher expression in CT than CS under cold condition are not related with over-represented categories but will be discussed here based on their likely relation with low temperature response/tolerance, according to the literature: *glutathione peroxidase* and *metallothionein* (antioxidant system), *fatty acid desaturase* and *phosphatidylinositol transfer protein* (lipid metabolism), *Tetratricopeptide Repeat-Containing Protein* (*TTL1)* (stress response). Glutathione peroxidases are thiol peroxidases that catalyze the reduction of H_2_O_2_ to water, being induced in response to cold stress. Passaia et al. (2013) showed that silencing the cold-inducible *OsGPX3* gene impairs normal plant development and leads to a stress-induced morphogenic response via H_2_O_2_ accumulation, which leads to smaller roots. Such response is similar to what we found in the CS genotype under low temperature stress (reduced root growth and H_2_O_2_ accumulation – Figures 1 and 2). Recently, Hsu and Hsu (2019) tested eight rice cultivars under cold treatment and found that the cultivars with higher growth rate had lower levels of H_2_O_2_ in the roots when compared with the low growth rate cultivars.

Metallothioneins, cysteine-rich proteins ubiquitous in eukaryotic organisms, are known to be involved in developmental processes and stress response, including low temperature in *Cicer microphyllum*, a wild relative of cultivated chickpea (Singh et al. 2011). Polyunsaturated fatty acids (PUFAs) play an important role in cold tolerance, since membrane fluidity is important to sustain the functional activity of membrane proteins and the membranes themselves (Beney et al. 2001). Recently, Wang et al. (2019) over-expressed an ω-3 fatty acid desaturase from *Glycine max* (*GmFAD3A*) in rice, obtaining a significant improvement in cold tolerance and survival ratio. Also, rice mutants deficient in ω-3 fatty acid desaturase (*OsFAD8*) fail to acclimate to low temperature stress (Tovuu et al. 2016). It is well known that root hair growth requires extensive cell wall modification. *OsSEC14-NODULIN DOMAIN-CONTAINING PROTEIN 1* (*OsSNDP1*) gene, encoding a phosphatidylinositol transfer protein (PITP), promotes root hair elongation via phospholipid signaling and metabolism, suggesting that the mediation of these processes by PITP is required for root hair elongation in rice (Huang et al. 2013). Mutations in the Arabidopsis *Tetratricopeptide-Repeat Thioredoxin-like 1 (TTL1 -* At1g53300) gene cause reduced tolerance to NaCl and osmotic stress that is characterized by reduced root elongation (Rosado et al. 2006). Therefore, we hypothesize that the *TTL1* gene found in our work could be the rice functional ortholog of Arabidopsis *TTL1*, functioning in cold stress response and contributing to the cold tolerance and maintenance of root growth under cold condition presented by the CT genotype (Figure 1).

On the other hand, under cold condition the CS genotype showed higher expression of genes related to several categories: amino acid metabolism, antioxidant system, calcium-related, defense, DNA structure maintenance, hormone-related, protein degradation/modification, transcription factor, translation, transport, and unknown (Table 1). For this genotype, only the genes that based on the literature are involved with cold sensitivity will be discussed. Among the defense-related genes, we found *SRC2* gene (Supplementary Table 3). In Arabidopsis, low temperature treatment induces expression of *AtSRC2* gene in roots, which binds to AtRbohF to enhance the Ca^2+^-dependent ROS production (Kawarazaki et al. 2013). Therefore, we hypothesize that the cold-inducible protein SRC2 found in CS genotype can function in cold response by controlling the Ca^2+^-dependent ROS production, which agrees with the ROS accumulation seen in this genotype under cold condition (Figure 2). The CS genotype presented higher expression of only one gene related with growth, *OsRAA1* (*ROOT ARCHITECTURE ASSOCIATED1* – Supplementary Table 3), a GTP-binding protein. Overexpression of *OsRAA1* promotes abnormal cell division (blocked transition from metaphase to anaphase during mitosis) and inhibits the growth of primary roots in rice (Xu et al. 2010), what could explain, at least partially, the reduced root growth presented by this genotype under cold condition (Figure 1). As seen under control condition, the CS genotype presented higher expression of ethylene-related genes than CT, including *1-aminocyclopropane-1-carboxylate oxidase* and *ethylene-responsive transcription factor* (Supplementary Table 3). As previously discussed, high levels of ethylene can inhibit root growth (Qin and Huang, 2018), and therefore we believe the root growth inhibition seen on CS genotype under cold condition (Figure 1) can be related with an increased ethylene level. A cytokinin-related gene (*cytokinin-O-glucosyltransferase 2*) was also found more expressed in CS than CT. It is already known that decreased cytokinin levels in roots results in increased root growth and higher drought tolerance in barley (Ramireddy et al. 2018). Therefore, higher cytokinin levels in CS than CT under cold condition probably contributed to the reduced root growth.

### Concluding remarks

The genetic improvement of root system has been regarded as an important approach to enhance crop production (Kitomi et al. 2018). Here we present a list of differential expressed genes that might be involved with cold tolerance and sensibility in rice plants. The model in Figure 7 summarizes the molecular and physiological responses of CT and CS genotypes under control and cold conditions. According to our data, high expression of genes related with cell wall remodeling (*glycosyl hydrolase*, *cellulose synthase*, *glycosyl transferase*, *wall-associated kinase*, *glycine-rich cell wall structural protein*), cytoskeleton and growth (*microtubule-associated protein 70*, *kinesin motor domain containing protein*, *growth regulating factor protein*, *auxin-independent growth promoter protein*, *RopGEF7*), signaling (*receptor-like protein kinase*, *Rapid Alkalinization Factor 21)*), antioxidant system (*glutathione peroxidase*, *metallothionein*), lipid metabolism (*fatty acid desaturase* and *phosphatidylinositol transfer protein*), and stress response (*Tetratricopeptide Repeat-Containing Protein*) might contribute to rice cold tolerance. On the other hand, high expression of the genes *SRC2* (defense), *root architecture associated 1* (growth), *ACC oxidase*, *ethylene-responsive transcription factor*, and *cytokinin-O-glucosyltransferase 2* (hormone-related) seems to be related with cold sensibility. Understanding more about the processes taking place in roots of plants subjected to cold stress can therefore not only provide a clearer picture of this complex phenomenon, but also disclose important information that can be used to develop engineering strategies aimed at improving plant tolerance to low temperature stress.

**Figure 7.**
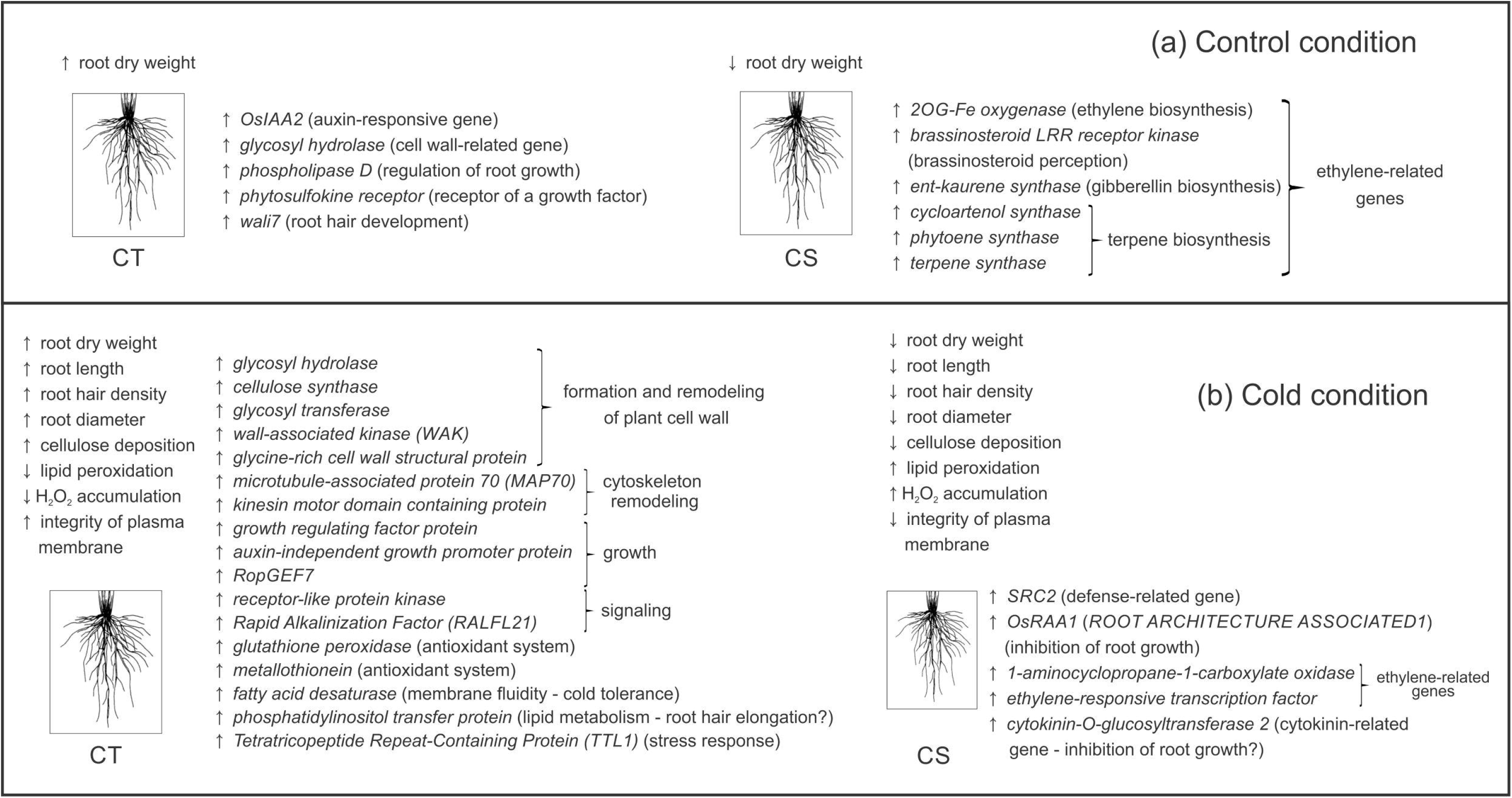
Gene expression profile and physiological responses of cold tolerant (CT) and cold sensitive (CS) genotypes under control (a) and cold (b) conditions.

CS: cold-sensitive
CT: cold-tolerant
DAB: diaminobenzidine
DEGs: differentially expressed genes
GA: gibberellin
QC: quiescent center

---

**Supplementary Figure 1.** Gene Ontology (GO) analysis of differentially expressed sequences from root transcriptomes of cold-tolerant (Tol) and cold-sensitive (Sen) genotypes under cold and control conditions through PlantGSEA (http://structuralbiology.cau.edu.cn/PlantGSEA/analysis.php). Terms enriched in either tolerant (black bars) or sensitive (white bars) datasets are shown as percentage of annotated genes in the dataset. All GO terms shown are differentially enriched in either group using Fisher’s Exact Test (p ≤ 0.05).

**Supplementary Table 1.** Gene-specific PCR primers used for RT-qPCR.

**Supplementary Table 2.** Differentially expressed sequences revealed by RNAseq in roots of cold-tolerant (Tol) and cold-sensitive (Sen) rice genotypes exposed to control (28°C) treatment for 24 h. Differential expression of the sequences marked with an asterisk were confirmed by RT-qPCR.

**Supplementary Table 3.** Differentially expressed sequences revealed by RNAseq in roots of cold-tolerant (Tol) and cold-sensitive (Sen) rice genotypes exposed to cold (10°C) treatment for 24 h. Differential expression of the sequences marked with an asterisk were confirmed by RT-qPCR.

**Supplementary Table 4.** Common genes present in the comparisons of cDNA libraries from roots of rice plants. Sequences more expressed in the cold-tolerant (CT) or cold-sensitive (CS) genotypes regardless the tested condition (control or cold treatment). Sequences marked in gray color presented opposite expression pattern in the CT and CS genotypes.

