## Supplementary figures and images for "Cold tolerance in rice plants is partially controlled by root responses"

### Supplementary Figure 1

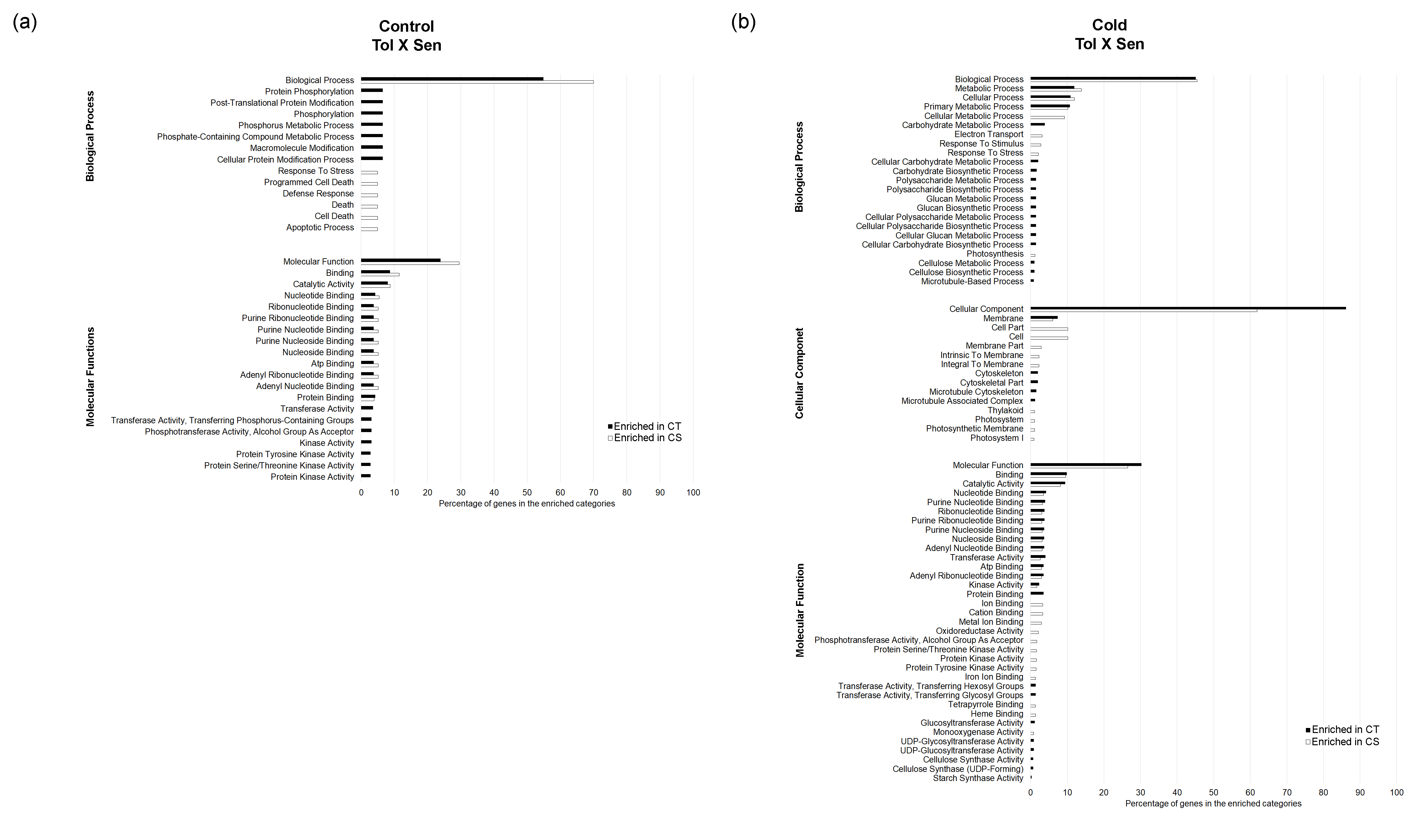
