## Supplementary Table 1 for "Cold tolerance in rice plants is partially controlled by root responses"

**Supplementary Table 1.** Gene-specific PCR primers used for RT-qPCR.

| **Gene ID (Phytozome**  **LOC_Os)** | **Gene** | **Forward primer**  **5’ → 3’** | **Reverse primer**  **5’ → 3’** | **Amplicon size**  **(bp)** |
| --- | --- | --- | --- | --- |
| LOC_Os04g02490 | *Expressed protein* | ATCTTGTTTCGCAGGATTGC | GAACGGCACAAATTGAAGGT | 134 |
| LOC_Os04g01330 | *Expressed protein* | CTTCCCCTCCTGTGAAAATG | CCTGTCATCAGATGGAATGGA | 81 |
| LOC_Os07g26100 | *Expressed protein* | AACCTTGTTTCGCAGGATTG | GAACGGCACAAATTGAAGGT | 134 |
| LOC_Os11g09850 | *Expressed protein* | CCACCTTGTGGTGTGAACTG | CAGAGTGCCATGTTTGCATC | 122 |
| LOC_Os07g12540 | *Expressed protein* | TTTGGTTGTGGGGAATTGAT | CCCTGGAACCAGGTAGACAA | 111 |
| LOC_Os07g20164 | *Expressed protein* | GCGTGTCTTTGGGTAGGTGT | GAAAGCGGGAGAGGGTTATC | 84 |
| LOC_Os04g02110 | *Disease resistance protein RGA3* | TAGAAGCAATGGGAGCGATT | CCATACGGGCTAACGACCTA | 142 |
| LOC_Os07g15440 | *Alanyl-tRNA synthetase family protein* | TGGTGGAGCCTCTGTAGTGA | CCACTGTTGCAAAAGAAAAACC | 138 |
| LOC_Os09g25934 | *Expressed protein* | ATCGCAGATGGATCCAAAAC | TCCCAGATGCTGAACAATCA | 140 |
| LOC_Os09g26554 | *Expressed protein* | GGAAGCAGCTGGTGTTGATT | AAGGCCCCTCATCATCTTCT | 132 |
| LOC_Os01g22490 | *OsUBQ5* | AACCAGCTGAGGCCCAAGA | ACGATTGATTTAACCAGTCCATGA |  |
